# Ultra-high temporal resolution 4D angiography using arterial spin labeling with subspace reconstruction

**DOI:** 10.1101/2024.07.03.601977

**Authors:** Qijia Shen, Wenchuan Wu, Mark Chiew, Yang Ji, Joseph G. Woods, Thomas W. Okell

## Abstract

**Purpose:** To achieve ultra-high temporal resolution non-contrast 4D angiography with improved spatiotemporal fidelity.

**Methods:** Continuous data acquisition using 3D golden-angle sampling following an arterial spin labeling preparation allows for flexibly reconstructing 4D dynamic angiograms at arbitrary temporal resolutions. However, k-space data is often temporally “binned” before image reconstruction, negatively affecting spatiotemporal fidelity and limiting temporal resolution. In this work, a subspace was extracted by linearly compressing a dictionary constructed from simulated curves of an angiographic kinetic model. The subspace was used to represent and reconstruct the voxelwise signal timecourse at the same temporal resolution as the data acquisition without temporal binning. Physiological parameters were estimated from the resulting images using a Bayesian fitting approach. A group of 8 healthy subjects were scanned and the in-vivo results reconstructed by different methods were compared. Due to the difficulty of obtaining ground truth 4D angiograms with ultra-high temporal resolution, the in-vivo results were cross-validated with numerical simulations.

**Results:** The proposed method enables 4D time-resolved angiography with much higher temporal resolution (14.7 ms) than previously reported (∼50 ms) whilst maintaining high spatial resolution (1.1 mm isotropic). Blood flow dynamics were depicted in greater detail, thin vessel visibility was improved, and the estimated physiological parameters also exhibited more realistic spatial patterns with the proposed method.

**Conclusion:** Incorporating a subspace compressed kinetic model into the reconstruction of 4D ASL angiograms notably improved the temporal resolution and spatiotemporal fidelity, which was subsequently beneficial for accurate physiological modeling.

## Introduction

Visualizing blood flow through the vessels in the brain provides valuable information for both diagnosis of many cerebrovascular diseases (e.g., stroke^1^, arteriovenous malformation ^2^, and steno-occlusive disease^3^) and surgical planning^4^. X-ray based methods (e.g., digital subtraction angiography) are widely used for assessment of vascular disease, yet exposure to radiation is known to carry some risk. Although contrast based MRI methods are radiation free, there is some concern about the use of gadolinium contrast agent exposure and accummulation^5–7^. Being one of the standard methods for neurovascular imaging, time-of-flight (TOF) MR angiography (MRA) acquires high spatial resolution blood vessel images without contrast agent. However, it is unable to provide hemodynamic information (e.g. to separate arterial and venous flow through an arteriovenous malformation), provides poor depiction of small and slow-flowing vessels and can suffer from venous contamination^8^.

Another non-contrast based method is arterial spin labeling (ASL)^9,10^ which labels inflowing arterial blood by inverting the magnetization at a labeling plane. Taking the difference between images with and without labeling provides information about blood flowing in the arteries or into the tissue, depending on the post labeling delay prior to image acquisition. Since static tissue signal is subtracted out, ASL images have zero background signal, which may enhance the visibility of small distal vessels^11^.

Further adding the information of how blood flow varies over time resulted in dynamic MRA which provides much richer information for clinical diagnosis^12^, and allows for the fitting of physiological models, providing quantitative metrics which could be sensitive to pathology^13^. Numerous methods have been developed recently to reconstruct time-resolved ASL angiograms. Some methods seek to encode temporal information into the ASL preparation process. Based on a Hadamard encoding scheme, the PCASL^9^ pulse train can be separated into blocks with or without labeling at each repeat^14,15^, followed by only one readout^16^ or a limited number readouts^17,18^. The time-resolved angiography information can then be reconstructed by “decoding” the resulting encoded images. Alternatively, a long train of temporally resolved readouts can be acquired immediately after each fixed labeling module to generate dynamic images^11,19,20^. High undersampling factors for each temporal frame is used to achieve high temporal resolution in short scan times. Advanced reconstruction methods can be adopted to leverage the complementary information available across multiple timepoints to mitigate undersampling artifacts. Song et al.,^21^ proposed K-space Weighted Image Contrast (KWIC)^21^ with radial sampling to reconstruct 4D dynamic MRA, which combined segmented spokes from different echoes at different parts of k-space. KWIC was later adopted in different dynamic MRA sequences^20,22^. Similarly, the keyhole method has also been exploited for time-resolved MRA^23^ reconstruction to combine data with different contrast while reducing artifacts.

Despite sophisticated design for data combination, these methods averaged readouts acquired at different timepoints with different contrast to achieve sufficient k-space sampling. They not only limited the temporal resolution that dynamic MRA could achieve, but also affected the spatiotemporal fidelity of the reconstructed images, since different k-space samples are acquired with the labeled blood at different spatial locations, especially for vessels with fast transiting blood. Inevitably, obtaining dynamic MRA without averaging data across timepoints would dramatically increase the undersampling factor. To compensate for this, improved priors for the MR signal would need to be incorporated to improve the conditioning of the reconstruction problem.

In this work, we aim to reconstruct ultra-high temporal resolution 4D ASL angiography images. Firstly, a kinetic model^13^ describing the angiographic signal during the course of a series of readouts after each ASL preparation was introduced to simulate a set of physiologically plausible signal curves. Secondly, the subspace technique developed by Tamir et al.^24^ was adopted to linearly compress this model and convert the poorly conditioned problem of reconstructing each time point into a better conditioned problem of estimating a small number of coefficient maps of these subspace components, greatly reducing the number of unknown variables and computational requirements. Thirdly, as well as conventional 3D radial sampling, a previously developed cone trajectory^25^ was integrated for efficient sampling of high spatial frequencies. Numerical simulations were performed to examine the spatiotemporal characteristics of our reconstruction and optimize the regularization weight. The in-vivo data was acquired with a combined angiographic and perfusion sequence (CAPRIA)^26^ but with only the angiogram reconstructed here. Altogether, high spatial resolution dynamic MRA images with temporal resolution equal to the sequence TR (∼15 ms) were reconstructed. The physiological parameters estimated from the in-vivo results were cross-validated with the numerical simulation for confirmation of the improvement in physiological modeling accuracy.

## Method

### Pulse sequence design

The pulse sequence used in this study was plotted in Figure 1. The sequence is composed of multiple repeats where an identical PCASL preparation is used but different k-space information is acquired. Within each repeat, a PCASL labeling or control module is followed by a single inversion pulse, as used in previous work^27^ to help suppress background tissue signal, before a series of readouts are acquired. The direction of each readout spoke followed the golden ratio^28^, the indices of which increased along the repeat dimension first as proposed by Song et al.^20^. Variable flip angles were applied to the train of readouts within each repeat to reduce the signal attenuation from the initial excitation pulses. In this work, two protocols were compared: one with a 3D radial trajectory and another with an in-house designed efficient cone trajectory^25^. The sequence design followed the combined angiographic and perfusion sequence^26^ (CAPRIA) although here we only focus on the reconstruction of angiographic images.

**Figure 1.**
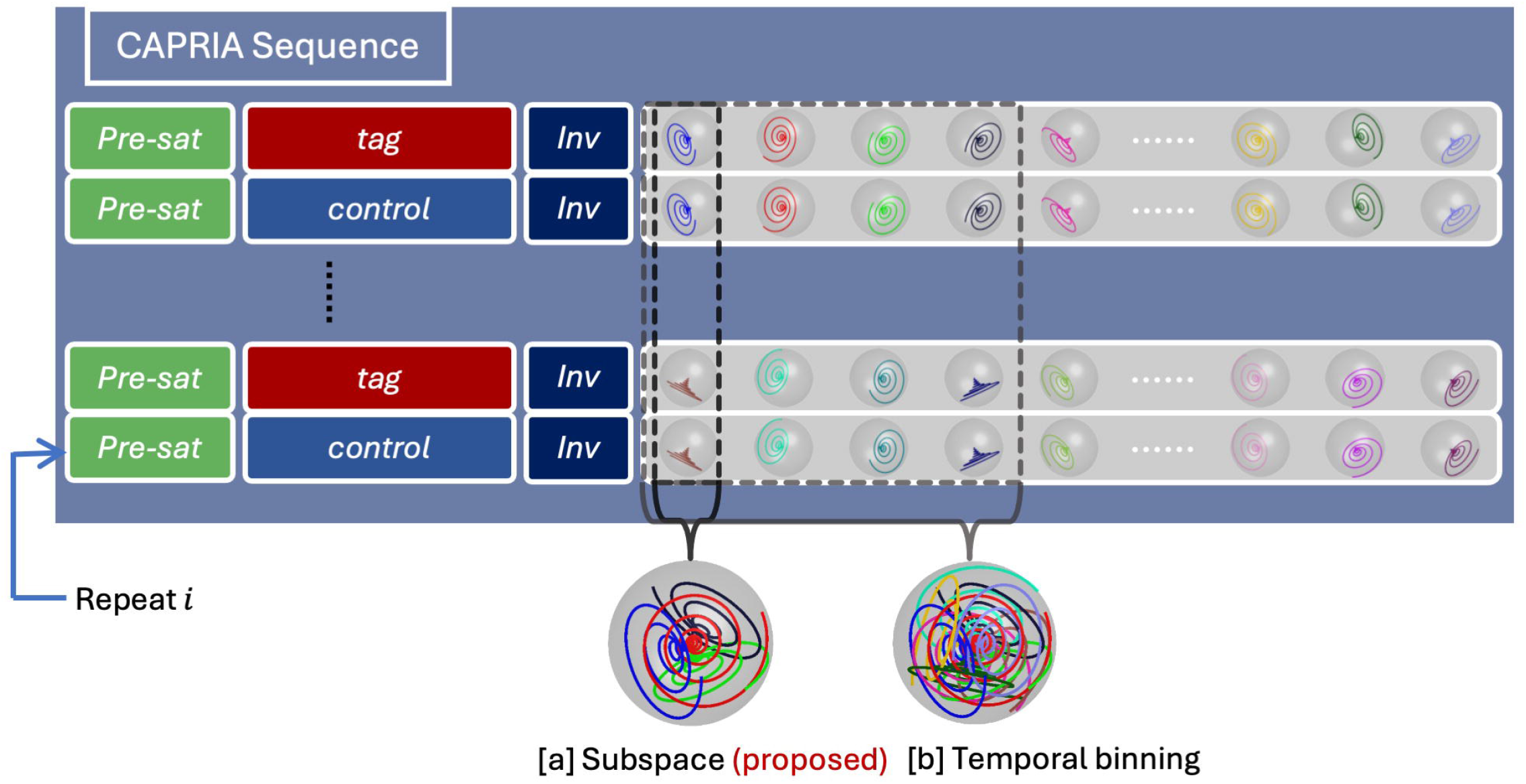
Overview of the CAPRIA pulse sequence with two different reconstruction methods: a) In the proposed subspace method, each temporal frame contains only readouts at a single timepoint. b) In contrast, in the temporal binning reconstruction scheme, each angiographic frame consists of readouts from several adjacent timepoints. Note that in the radial protocol, each cone trajectory plotted in the figure would be replaced by a radial spoke aligned with its central axis.

To reconstruct a dynamic angiogram from the acquired data, conventionally, readouts from multiple TRs would be grouped to achieve sufficient k-space sampling, as shown in Figure 1b, which we term the “temporal binning” method. However, the proposed method, referred to as the “subspace” method in Figure 1a, combined spokes within only one TR for each frame, achieving a temporal resolution same as the TR. Further implementation details are explained below.

### Angiographic kinetic model

Given the sequence parameters (i.e., labeling duration, TR, flip angle, etc.), the angiographic signal evolution within each voxel after PCASL labeling was explicitly modeled by Okell et al.^13^, with 4 independent parameters: the blood transit time from the labeling plane to the imaging voxel, *δ*_*t*_; the sharpness, *S*; and time-to-peak, *p*, of a gamma variate dispersion kernel, and a scaling factor proportional to blood volume, *A*. The model describing the signal variation across time is composed of effects from dispersion, relaxation and attenuation due to RF pulses, which are explained below.

*D*(*t*_*d*_): The ideal temporal profile of the blood bolus is a rectangular function with a width equal to the labeling duration. Depending on the dispersion time-to-peak and sharpness, this ideal profile is blurred, which is described by its convolution with a gamma variate kernel, *D*(*t*_*d*_), while flowing through the vascular tree on the way to the voxel of interest, as described by Equation 1. *t*_*d*_ is the transit time delay due to blood dispersion.

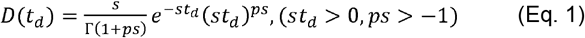

*T*(*δt,t*_*d*_): Additionally, the inverted magnetization of blood experiences T_1_ recovery as described by *T*(*δt,t*_*d*_) in Equation 2.*t*_1*b*_ is the T_1_ of arterial blood.

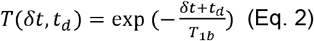

*R*(*t*_*d*_):The attenuation of the longitudinal magnetization difference signal by the imaging RF pulses is described by *R*(*t*_*d*_)in Equation 3, making the simplifying assumption for this 3D acquisition that all labeled blood experiences all RF excitation pulses before *t*_*d*_. *TR* is time for one RF pulse to the next in the train of readouts.

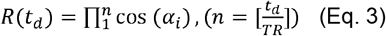

*S*(*t*): Finally, the actual signal amplitude depends on the flip angle at each excitation, contributing an additional sin*(α*_*i*_) term, as described previously^26^. Combining all these effects, the final signal profile *S* at each time after labeling *t* and each voxel position is characterized in Equation 4.

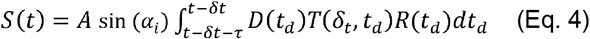

### Subspace generation

The prior knowledge of the signal time series was incorporated into the reconstruction using a subspace method. Specifically, a plausible range of model parameters was chosen as follows: *δ*_*t*_: 0.1 ∼ 2.0 (*second*), *p*:0.001 ∼ 0.5 (*second*), *s*;1 ∼ 20 (*second* ^− 1^) (Note the parameter name s is different from second as a unit). To cover a wide range of temporal variations for blood flow, 40 equispaced samples were drawn for each parameter within the range and combined, forming a dictionary with 40^3^ samples in total. The scaling factor *A* was not included in the dictionary generation process as it would not affect the subspace. As these different signal time courses are strongly correlated with each other, they lie on a low-dimensional manifold embedded in a high-dimensional space. Although the exact manifold is not easy to find, a linear subspace can serve as a sufficient approximation, with straightforward calculation and computational convenience^24^. Principal component analysis (PCA) was applied to the generated dictionary, as shown in Figure 2a. The space spanned by a small number of principal components was regarded as the subspace for the signal variation. As plotted in Figure 2b, the first few principal components capture slow signal variations, with higher order components containing more rapid oscillatory behavior. Using more principal components can give a better approximation to the signal variation but requires higher computational cost. To determine the number of components required to strike a good balance between computational efficiency and accuracy, the relative error 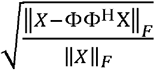 was calculated on the whole dictionary as shown in Figure 2c. *X* is the generated dictionary and Φ is the subspace. We found that using the first 12 components resulted in 0.961% relative error while keeping the same size of unknown parameters to solve as in reconstructing 12 temporally-binned frames^25,26^. The NRMSE at each timepoint is also shown in Figure 2d. Using 12 principal components achieved greatly reduced oscillatory errors compared to 4 or 8 components. Further increasing the number of components from 12 to 16 did not bring about significant improvements but increased computational requirements. Therefore 12 principal components were chosen for the subspace reconstruction for angiography in this work.

**Figure 2.**
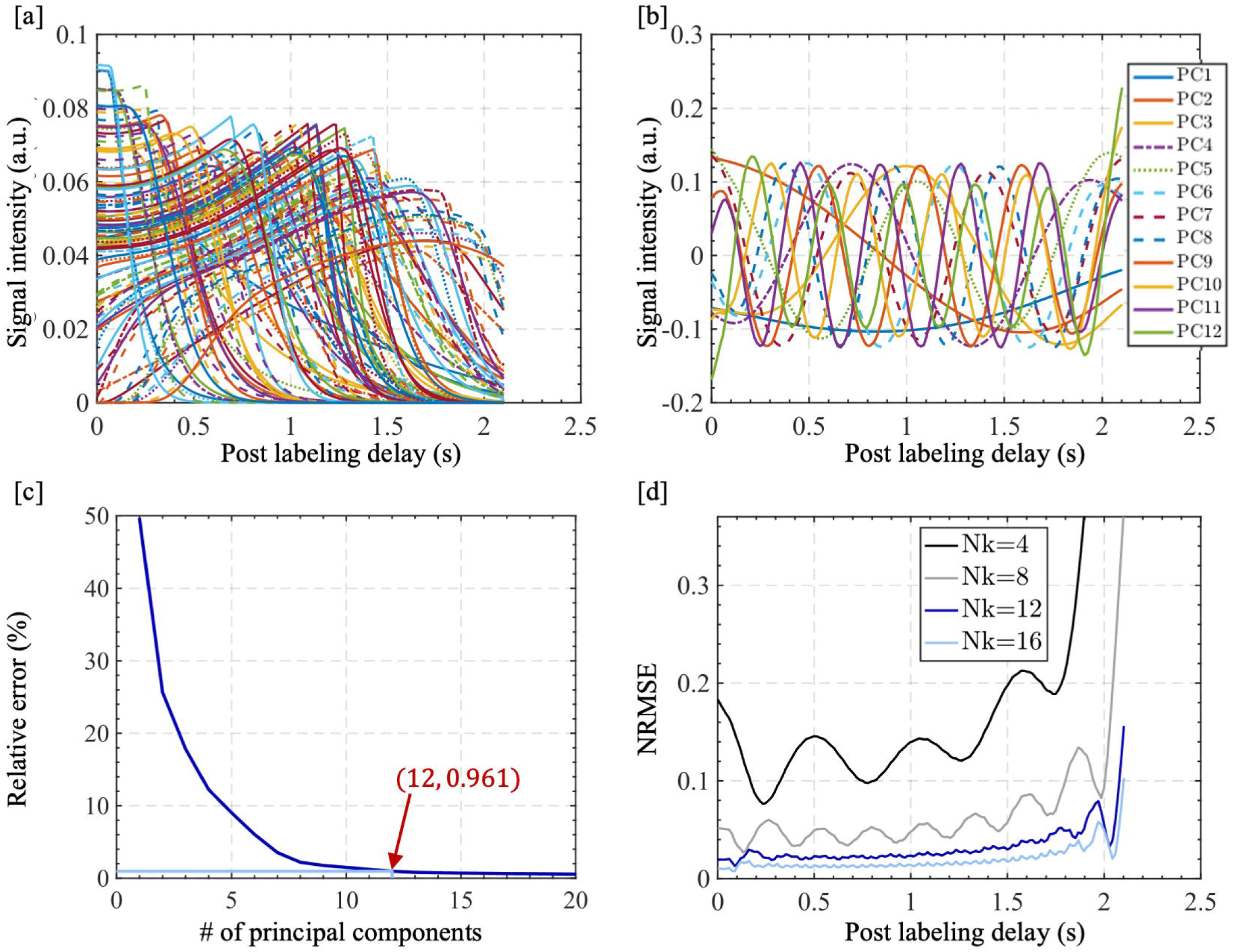
Subspace generation for the angiographic time series. a) Simulated dictionary of angiographic time series across different combinations of plausible physiological parameters. b) The first 12 principal components (PC) were extracted using PCA. Curves of PCs with larger indices manifest higher oscillating frequency. c) Relative reconstruction error of the dictionary with different number of principal components used for angiography signal compression. d) NRMSE at each timepoint with a different number (Nk) of principal components.

### Subspace reconstruction for angiographic images

The target time series, *X*, can be approximated by the weighted combination of principal components, Φ. As shown in Equation 5, the voxel-wise weights for each principal component, *α*, are to be estimated. The reconstruction problem was converted to the form of Equation 6, where *W*_*i*_ was the *i*^th^ patch extraction operator, *p* was the sampling mask, *F* was the Fourier transform and *C* was the coil sensitivity. Similar to previous work^24^, locally low rank regularization^29^ (LLR) was applied across the principal component coefficient maps to reduce noise. *λ* was the weight adjusting the LLR regularization.

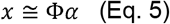

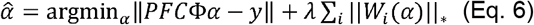

As operators *F,C* were applied on the spatial dimension, independent from the operator Φ on the temporal dimension, the original order of operators could be modified to *P*Φ*FCα*. The combined operator Φ^H^*P* Φused in every iteration could be calculated before the start of iterative optimization, so that the whole optimization process could be performed entirely in the subspace domain without expanding to a full time-series^24^. The technique reduced the computation time of subspace reconstruction to that of the temporal binning reconstruction.

### Temporal binning reconstruction method

The proposed subspace reconstruction was compared against the method used to reconstruct 4D dynamic MRA in previous work^25,26^. As presented in Equation 7, the LLR regularization was directly applied to the dynamic image *x*. Each frame involves grouped k-space data sampled from multiple TRs. Note the meaning of each notation was shared between Equations 6 and 7.

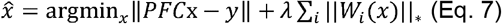

### Numerical simulation

Before in-vivo acquisition, numerical simulations were performed to examine the spatiotemporal characteristics of the subspace reconstruction results compared to the original temporal binning reconstruction approach. A line with one voxel width crossing the center of every 3D volume and varying across time was created, simulating a time-series of a single vessel. The four parameters of the kinetic model in Equation 4 (i.e., *δ*_*t*_,*p,s,A*) were distributed along the line in a linearly varying fashion, simulating increasing transit time and greater dispersion along the length of a vessel. Following the previous literature^13^, *δ*_*t*_ ranged between 0.25 and 1.8s, *p* ranged between 0.1 and 0.5*s, s* ranged between 1 and 10_*s*_ ^−1^, and the scaling parameter *A* was set to be equal to 1 along the simulated vessel. A 4D time-series with the same spatial resolution (1.1 *mm*) as in-vivo reconstruction was simulated with these parameters and the kinetic model. The TR, number of readouts and k-space trajectories in the simulation were also kept the same as for the in-vivo acquisition protocols. After transforming the 4D timeseries to k-space with a single coil and uniform sensitivity using the forward NUFFT^30,31^, temporal binning and subspace methods were used to separately reconstruct from the k-space data. In order to compare the spatial resolution, a point spread function was also calculated along the direction perpendicular to the simulated line. Both the radial trajectory used in Okell et.al.,^26,27^ and the cone trajectory used in Shen et al.^25^ were simulated and reconstructed with the two methods. A variational Bayesian method (Fabber^32^, part of FSL^33^) was used to infer the four model parameters from the reconstructed images and evaluate the reconstruction quality. Additionally, weights 1e-4, 5e-4, 1e-3, and 1e-2 for the LLR regularization term were tested for subspace reconstruction to inform the in-vivo scenario. The data transformed from numerical phantom to k-space was scaled in the same way as described in in-vivo scenario later before iterative reconstruction to ensure the regularization weight transferrable to in-vivo reconstruction. To align with the optimal design in previous literature^25,26^, the regularization weight of 1e-1 was used for temporal binning reconstruction.

### In-vivo acquisition and reconstruction protocols

To evaluate the proposed reconstruction method, a group of 8 subjects were scanned on a 3T Prisma scanner (Siemens Healthineers, Erlangen, Germany) using a 32-channel receive-only head coil under a technical development protocol agreed by local ethics and institutional committees, and they all provided consent.

For each subject, two back-to-back scans with two different protocols were acquired. The two protocols followed the same design as the “cone protocol” and “matched radial protocol” in the previous literature^25^. The variable flip angles varied quadratically across the readout and ranged between 3°-12°. The TR for each readout was 14.7 ms, where the readout time was 10 ms. Note that the ASL labeling duration (1.8 s), the total time for all readouts after each ASL labeling module (2116.8 ms) as well as the total scan time (6 min 12 s) were kept consistent between the two protocols. The FOV was 200 × 200 × 125 *mm* and the spatial resolution was 1.13 *mm* isotropically.

Before reconstruction, the original 32-channel data were compressed to 8 virtual coils^34^. Coil sensitivity map estimation used the adaptive combination method^35^. For the temporal binning reconstruction, we followed previous literature^25^ and used 1e-1 as the LLR regularization weight. For the subspace reconstruction, differences in data scaling between simulated and acquired data were removed by dividing both k-space data sets by their 95-percentile magnitude value before reconstruction to allow transfer of the finetuned regularization weight from numerical simulation to in vivo data. Based on the numerical simulation, a regularization weight of 5e-4 was adopted for subspace reconstruction.

Apart from the subspace and temporal binning reconstruction methods mentioned above, two advanced image reconstruction methods were also included for comparison, where the temporal resolution was matched to that of the subspace method. Firstly, we adapted the LLR regularized reconstruction used by temporal binning to achieve the same temporal resolution as the subspace method, termed “LLR-matched” below. Another method “Extreme MRI”^36^ which combined multi-scale low rank and stochastic reconstruction was also used for comparison. For both methods, different regularization weights were investigated, the details of which are shown in Supporting Figures S1 and S2.

## Results

### Numerical simulations

Simulated results on a single-vessel numerical phantom are shown in Figure 3. In the 4D phantom (Figure 3a), the blood bolus flows from left to right, with early blood arrival and minimal dispersion on the left and delayed arrival and greater dispersion moving to the right. The signal time-series from three representative locations along the simulated vessel are displayed to compare the reconstruction fidelity between the subspace and temporal binning methods. Where the ground truth signal varies rapidly on the left of the phantom (early arrival and low dispersion), the subspace reconstruction cannot fully represent this rapid signal change and there is a slight discrepancy between the reconstructed signal and the “ground truth”. However, the subspace method still captures the overall signal trend. In contrast, the temporal binning signal curve drops from the beginning and fails to delineate the plateau and sharp transition because of averaging data points across a wide temporal window. Towards the middle and right of the phantom where the signal arrives later and is more dispersed, the subspace method gives a very good approximation to the “ground truth” signal, whereas the temporal binning result still deviates from the “ground truth” before the signal peak. Additionally, in the temporal binning results, when the peak time lies between two reconstructed frames the resulting signal curve shows a plateau rather than a single peak, as shown in Figure 3a.

**Figure 3.**
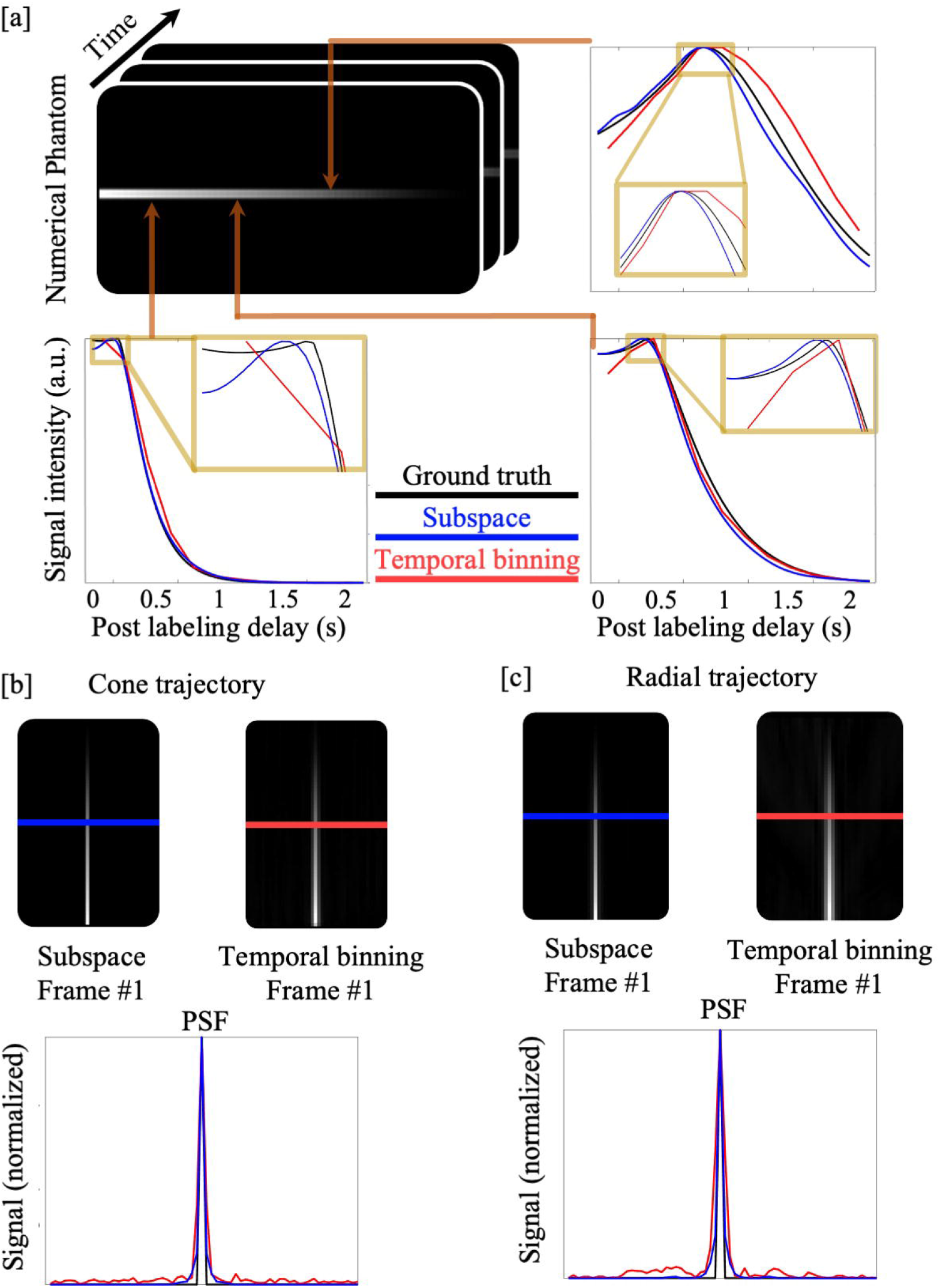
Spatiotemporal comparison of simulated results between the subspace and temporal binning reconstruction methods. a) The signal timecourses sampled from three points along the simulated vessel are shown. The curves reconstructed using the subspace method approximated the “ground truth” with better accuracy. b,c) The point spread function from the plane perpendicular to the simulated vessel using both cone (b) and radial (c) trajectories for the two reconstruction methods. Combined with cone and radial trajectories, the subspace method showed a thinner central lobe, lower sidelobe aliasing, and improved delineation of this narrow structure than temporal binning in both cases.

In Figure 3b-c, the spatial characteristics of both reconstruction methods as well as both sampling trajectories were examined. Since the phantom was one voxel wide, the point spread function (PSF) on the plane through the volume center and perpendicular to the line was plotted. In Figure 3b-c, regardless of the sampling trajectory, the central peak was thinner in the subspace PSF than it was for temporal binning. The sidelobe aliasing was also better suppressed when using the subspace reconstruction approach compared to the original temporal binning method for both cone and radial trajectories. Figure 4 shows how the improvement in the temporal fidelity of the reconstructed signal with the subspace approach translated into improved angiography parameter estimation. In Figure 4a, parameters estimated with the temporal binning method exhibited a discrete and stepwise pattern, unlike the gradual variation in the ground truth image. As shown in the zoomed-in region in Figure 4a, one cause of these errors was when the true signal peak was not captured in the temporal binning reconstruction due to the limited temporal resolution. In comparison, the parameters estimated from the subspace approach were smoothly varying and closer to the ground truth. In Figure 4b, the relative error of the temporal binning and subspace methods with different regularization weights were plotted. It could be observed that parameters of temporal binning were mostly overestimated and deviated from the ground truth much further than the subspace results. In addition, the most accurate quantification was achieved when the regularization weight was set to 5e-4 in the subspace reconstruction, which was adopted for subspace reconstructions of the in-vivo data below.

**Figure 4.**
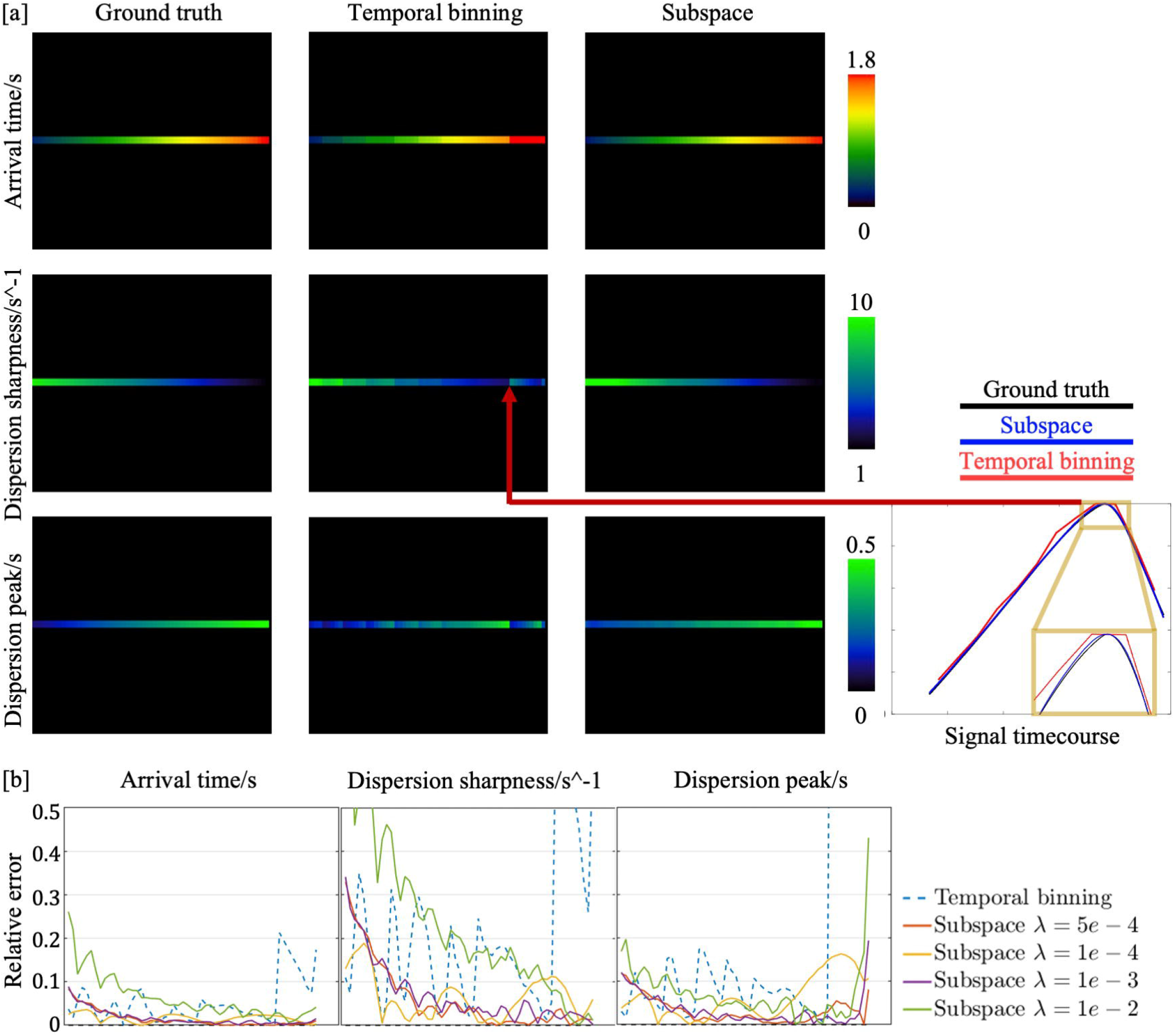
Parameter estimation from numerical phantom images reconstructed by subspace and temporal binning methods using the cone trajectory. a) Comparison of estimated dispersion parameters and arrival time between the two methods. b) Relative error of the estimated parameters along the simulated line compared to the ground truth. Parameters estimated from the subspace method with different regularization weights were explored.

### In-vivo angiography results

In-vivo angiograms reconstructed with subspace and temporal binning methods are compared in Figures 5 and 6. Note that the vasculature was already filled with blood at PLD=88ms because a relatively long (1.8s) labeling duration was used in the protocol.

**Figure 5.**
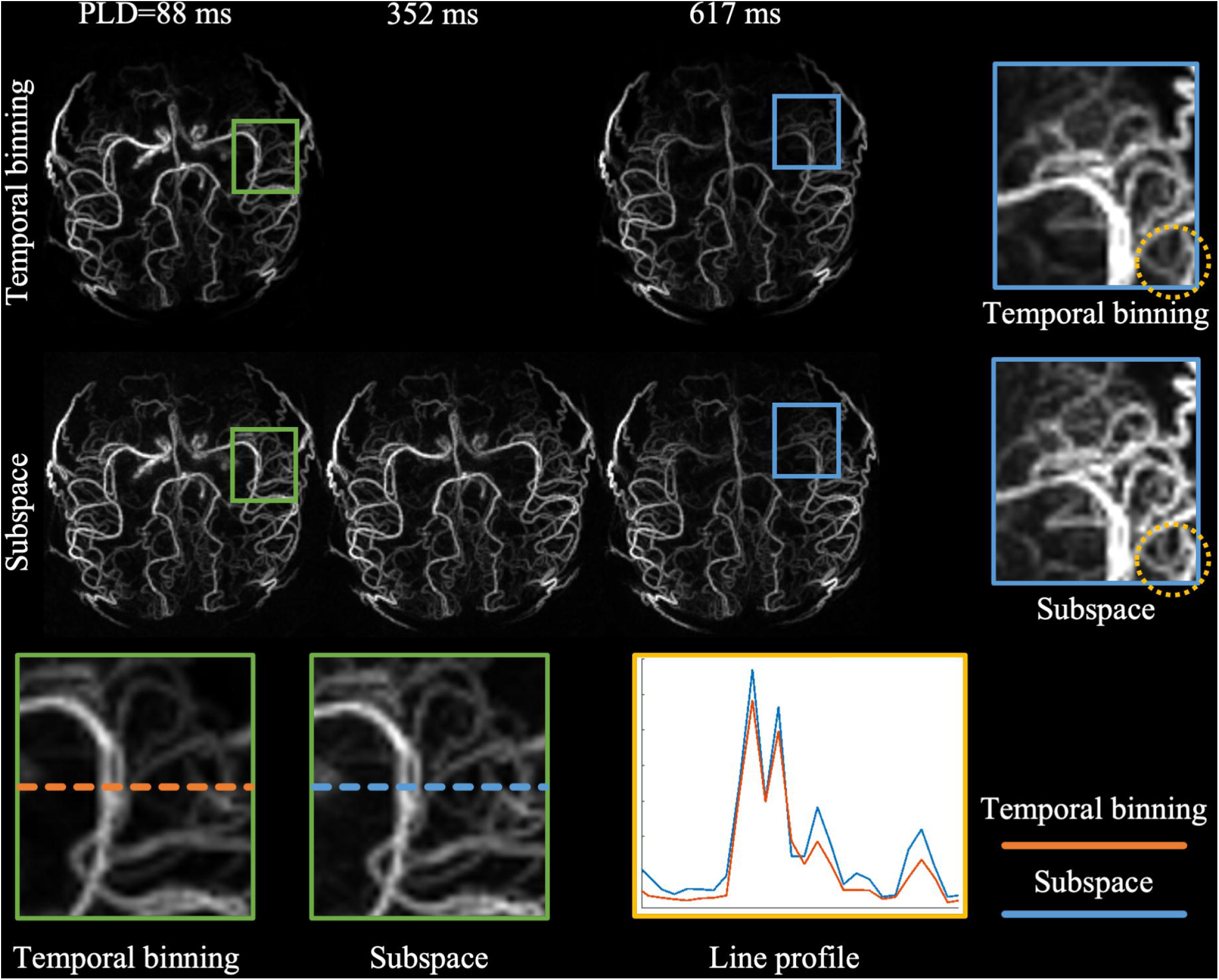
Spatial comparison of dynamic MRA acquired with a radial trajectory and reconstructed by subspace and temporal binning methods. Note that the vasculature was already filled with blood at PLD=88ms as a relatively long labeling during (1.8s) was used. The quoted PLD represents the center of the temporal window for each frame. The temporal binning result did not have the image at PLD=352ms due to limited temporal resolution. Regions in the green box were zoomed for comparison of the visibility of thin vessels. Line profiles were sampled within the green box for both reconstruction methods. Regions enclosed in the blue box were also zoomed in to confirm the better visibility of vessels in the subspace results across time. The yellow circle marked an example where thin vessels were hard to distinguish from the background when using temporal binning but became much clearer when using the subspace approach.

**Figure 6.**
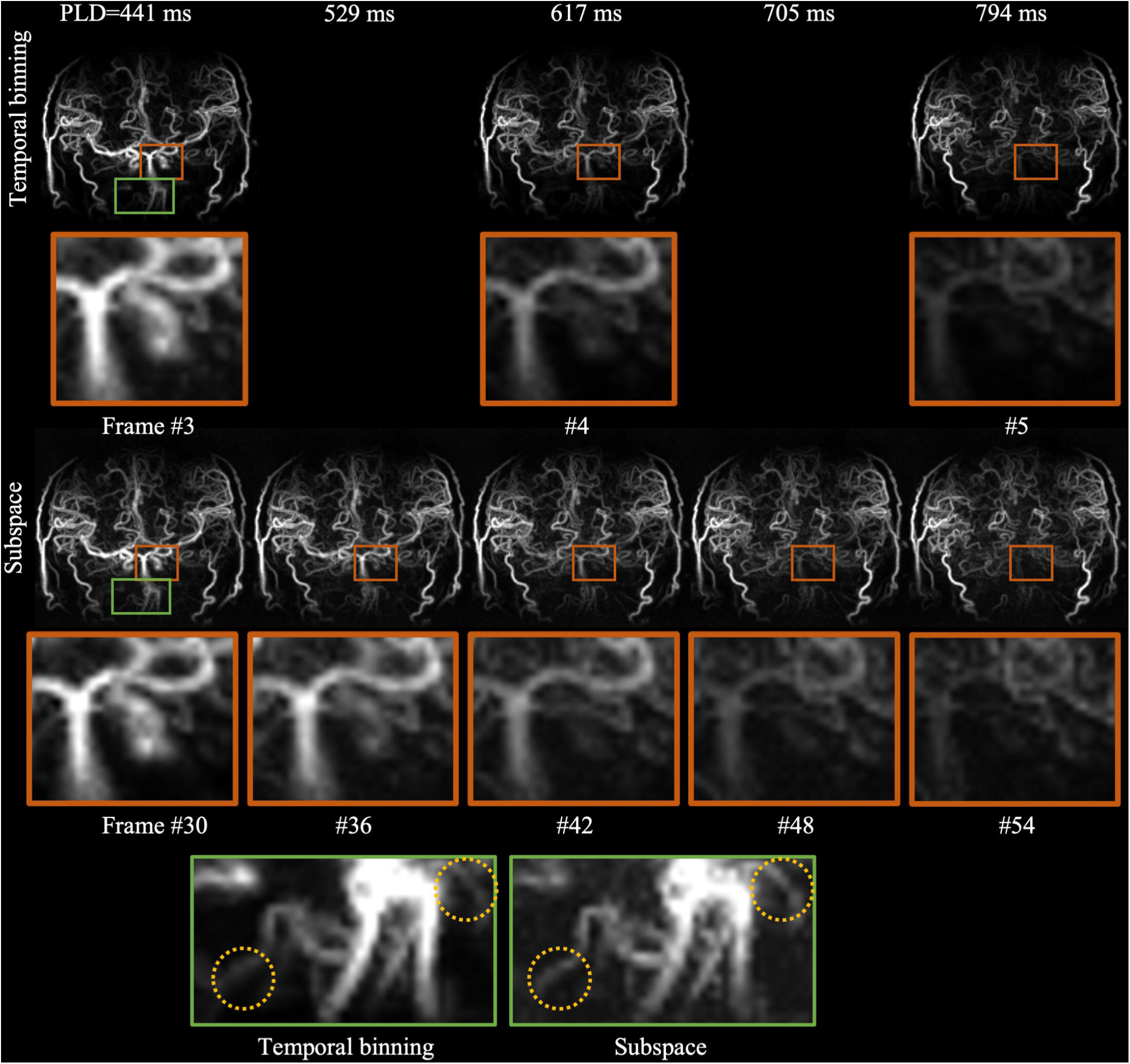
Comparison of in vivo dynamic MRA acquired with a cone trajectory and reconstructed by subspace and temporal binning methods at different post labeling delay times. 3 out of 12 frames reconstructed by temporal binning were shown on the top row. Similarly, among the 144 frames in total reconstructed by subspace method, 5 equispaced frames between 441 ms and 794 ms after labeling are also shown. 3 of them matched the timepoints of the consecutive frames #3∼5 of temporal binning result. The blood flow dynamics shown in the zoomed-in regions demonstrated superior temporal discernability of subspace method compared to temporal binning. The zoomed-in green box further showed evidence that the subspace method provided better clarity of vessels than temporal binning as highlighted by the yellow circles.

The notable improvement in spatial fidelity could be observed in images reconstructed from data acquired with a radial trajectory (Figure 5). Consistent with the simulated results in Figure 3b, the angiograms generated by the subspace method show improved delineation of fine vessels compared to temporal binning. For instance, more vessels could be visualized with better clarity in middle cerebral artery (MCA) in subspace result compared to temporal binning result as shown in the enlarged blue box in the figure. The relative signal intensity of vessels compared to background was further investigated with line profiles, where the peak to bottom difference were noticeably larger in subspace data, indicating that vessels were more discernable from the background in the subspace reconstruction.

In Figure 6, the data acquired with the cone trajectory was also reconstructed with temporal binning and subspace approaches respectively. Three consecutive frames in temporal binning angiography are shown in the first row. The subspace results have much higher temporal resolution than the temporal binning results. Between every two successive temporal binning frames, 10 more frames were reconstructed using the subspace method, demonstrating much smoother signal variations. Note, only one intermediate frame is shown here in Figure 6 for brevity. Regions around the internal carotid artery (ICA) were zoomed in from each frame and the dynamic of blood flow was better depicted temporally in subspace method than in the temporal binning method. In addition, the spatial fidelity of MRA was also improved in the subspace approach compared to temporal binning as illustrated in the zoomed-in green regions.

### Quantitative comparison

The time-resolved MRA in vivo data reconstructed by the two different methods was further analyzed to extract physiological parameter maps (Figure 7). In agreement with the simulation results in Figure 4a, the in vivo arrival time estimated from temporal binning exhibited stepwise discontinuities along the arteries (Figure 7a), while the subspace data gave a more realistic smooth, continuous increase in transit time along the arteries, thanks to its ultra-high temporal resolution. Additionally, although there is no ground truth for this in vivo data, the arrival time appears to be over-estimated with temporal binning when the transit times are close to the length of the readout train (∼2.1s). This overestimation was observed in both simulated and in-vivo results, exemplified in the white circle, but was noticeably improved in the subspace results. The blood dispersion parameters also benefited from the improved temporal resolution and fidelity available with the subspace method as shown in Figure 7b. For instance, the dispersion sharpness of temporal binning MRA in the distal arteries was much higher than with the subspace reconstruction, which is inconsistent with previous work^13^ and the expectation that dispersion will continue to increase as blood passes down the vascular tree. This observation was also consistent with the simulation results (yellow circle).

**Figure 7.**
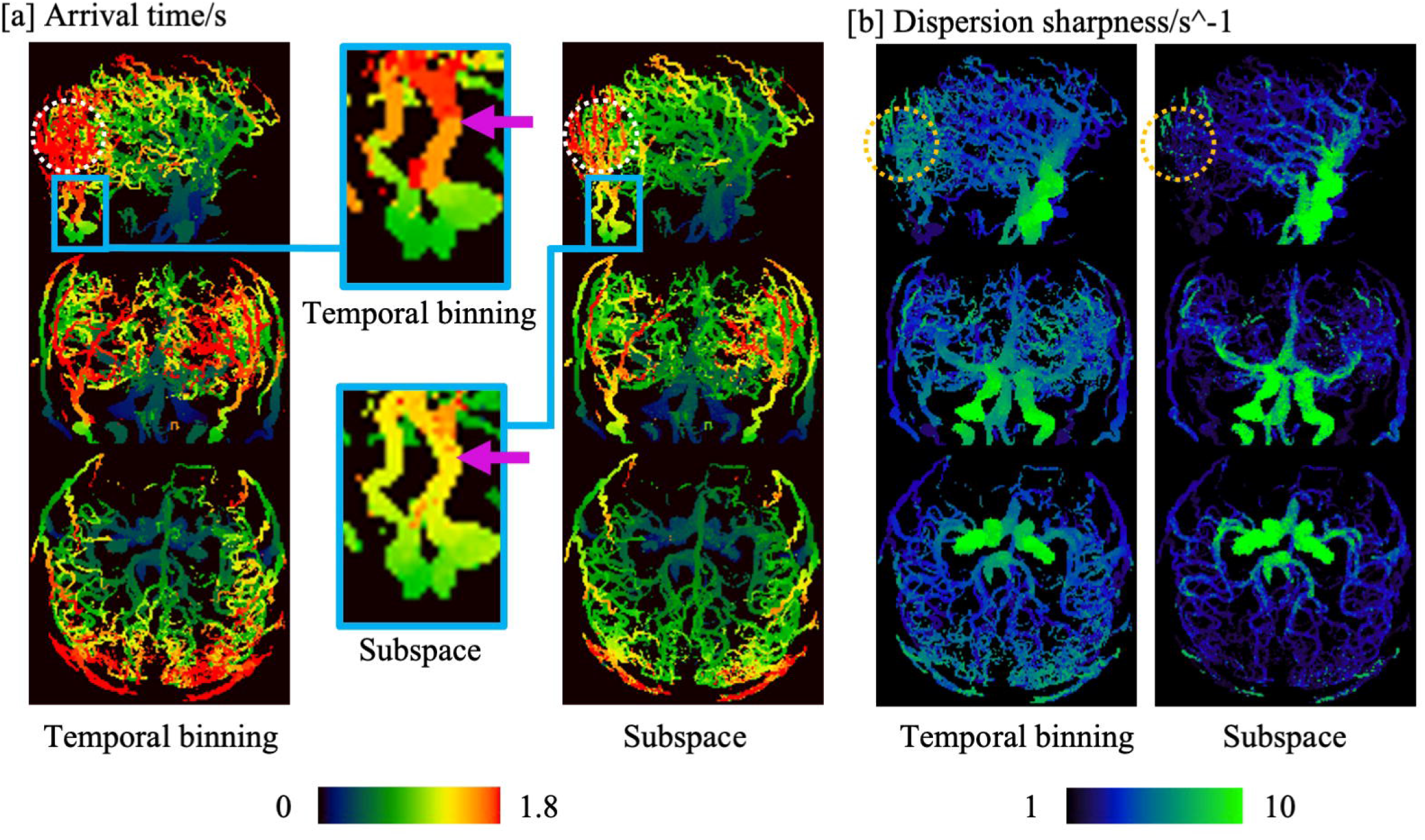
Comparison of blood arrival time and dispersion sharpness estimated from temporal binning and subspace results. a) Arrival time in temporal binning MRA was overestimated and discontinuous (highlighted by the purple arrows), whereas that in subspace was smoother and more realistic. b) The dispersion sharpness estimated from temporal binning also demonstrated an increase in very distal vessels, which would not be expected as dispersion should continue to increase along the path of the vessels, whereas this effect was much less apparent in the subspace data. These results are consistent with the numerical phantom simulations shown in Figure 4.

Parameter map histograms derived from the temporal binning and subspace reconstruction methods are shown in Figure 8, both for a single subject (Figure 8a) and for all subjects (Figure 8b). In both cases, the arrival time distribution for the temporal binning data showed discrete peaks, unlike the smoother distribution observed with the subspace data, most likely caused by the limited temporal resolution of the temporal binning data. As was observed in Figure 7, the dispersion sharpness appeared to be overestimated in temporal binning relative to the subspace approach. Although the dispersion peak was also likely biased in the temporal binning results based on our simulation results, the direction of deviation varied (Figure 4a).

**Figure 8.**
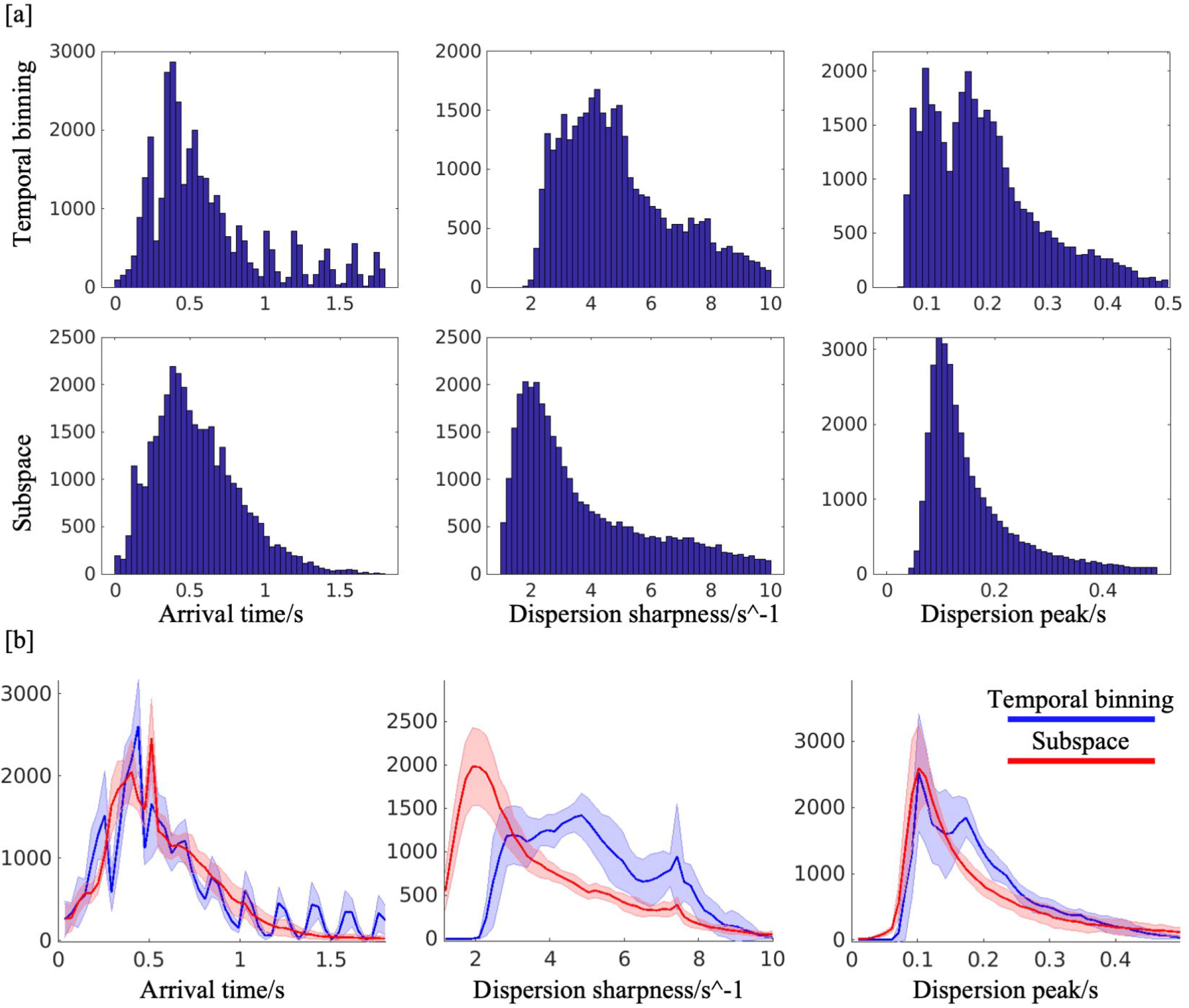
a) Histogram of three angiography parameters estimated from one subject. b) The distribution of these parameters on the group level. The average count of each value was plotted with the “red” and “blue” curves corresponding to subspace and temporal binning reconstruction methods, respectively. The standard deviation of the voxel count across subjects was added as margins around the curve.

### Comparison with other advanced reconstruction methods

In Figure 9, the angiography results were further compared against other advanced methods reconstructed with the same temporal resolution as the subspace method from the same k-space data. As all 144 frames were explicitly reconstructed, the “LLR-matched” approach had much higher computation time and memory requirements (300GB memory, 28hr reconstruction time) than the subspace method (130 GB memory, 4hr reconstruction time). In addition, since 12x more frames were reconstructed than temporal binning without introducing additional regularization, LLR-matched exhibited degraded image quality and only the large arteries with strong signal intensity could be visualized while the small distal vessels were lost as shown in the green box. The reduced sampling per frame of LLR-matched also increased its dependence on locally low rank regularization, which introduced artifacts in arteries highlighted by the red arrow. Extreme MRI used an even more sophisticated reconstruction with multi-scale low rank patches and resulted in better sharpness of small vessels than LLR-matched. However, SNR was low and image details were inferior to those of the subspace reconstruction as could be seen from the enlarged green box. Overall, the subspace results demonstrated the best temporal and spatial fidelity and resolution among these reconstruction methods.

**Figure 9.**
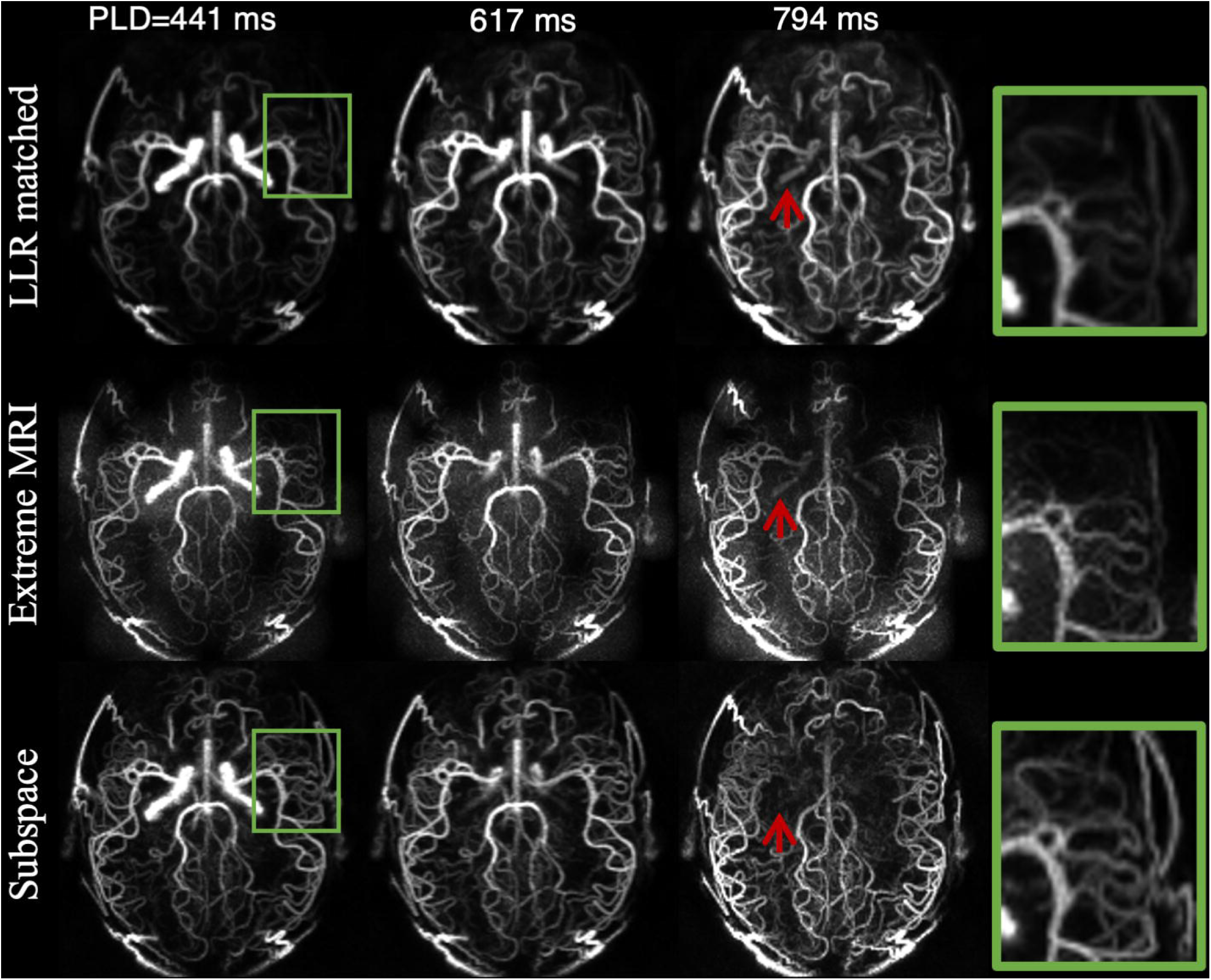
Comparison of subspace with other advanced reconstruction methods. LLR-matched adapted the temporal binning reconstruction to the same temporal resolution as subspace. Extreme MRI is a multi-scale low rank method and also had matched temporal resolution. Regions enlarged within the green box show loss of details in ExtremeMRI and LLR-matched results, but better clarity in subspace. Red arrows highlighted falsely reconstructed arteries in LLR-matched due to regularization. Further investigation of the effects of regularization weight in ExtremeMRI and LLR-matched is shown in Supporting Figures S1 and S2.

## Discussion

In this work, a subspace, generated by linearly compressing a kinetic model, was incorporated into the reconstruction of 4D dynamic angiograms to achieve ultra-high temporal resolution (14.7ms) and improved spatiotemporal fidelity. Our proposed subspace reconstruction was optimized through numerical simulations, where the optimal number of subspace principal components was chosen to balance the relative fitting error and computational requirements. Simulations were also used to find the optimal regularization weight, which is challenging to do using in-vivo data where there is no ground truth. The reconstructed in-vivo results were compared between the subspace and temporal binning methods and validated against the simulation results both qualitatively (improvement of spatiotemporal fidelity) and quantitatively (improvement of physiological modeling accuracy). Additionally, the proposed method was shown to outperform other advanced methods (“LLR-matched” and ExtremeMRI^36^) when the same spatial and temporal resolution were used.

Incorporation of the modality specific prior knowledge (i.e., the kinetic model) was essential in obtaining ultra-high temporal resolution 4D angiograms without increasing scanning time (∼6 min), as the high undersampling factor per frame presented a very ill-conditioned reconstruction problem. Taking advantage of compressed sensing^37^ (e.g., low rank regularization) and adjusting the regularization weight could alleviate the problem of insufficient k-space sampling while also potentially introducing artifacts. As shown in Supporting Figures S1 and S2, both LLR-matched and ExtremeMRI^36^ methods were evaluated with different regularization weights, and the results were either too noisy (i.e., under regularized) or too blurry (i.e., over regularized), with loss of distal vessel visibility. These results demonstrate that relying solely on locally low rank regularization limited the temporal variation to compensate for the insufficient sampling per frame and was not enough for faithful reconstruction of ultra-high temporal resolution 4D angiograms. In contrast, our subspace method incorporated a signal model of blood flow into the reconstruction, limiting the degrees of freedom. The fact that k-space samples acquired across the whole train of readouts can contribute to the estimation of each subspace coefficient map likely improves the spatial fidelity of the reconstruction. The subspace approach also avoids the temporal misallocation of signal which is inevitable in the temporal binning method. In our subspace method, a locally low rank regularization term was also included on the coefficient maps to reduce noise.

Several characteristics of the sequence used to acquire our data also contributed to achieving ultra-high temporal resolution 4D angiograms. Firstly, the 3D golden angle^28^ and repeat-first ordering^20^ in the sequence guaranteed similar and approximately uniform k-space sampling at each timepoint, supporting a frame window size of one TR in the subspace reconstruction. Voxels at different timepoints were also assumed to be spatially aligned for the kinetic model constraint to be applicable voxel-wise. As the total readout time within each repeat is ∼2s, it is unlikely that significant motion will occur within each train of readouts. Since the between-repeat motion was shared for samples acquired across all TR periods, motion would be more likely manifested as blurriness rather than spatial misalignment between TRs, which is expected to make the subspace constrained reconstruction of 4D angiograms less motion sensitive.

Additionally, the proposed reconstruction was particularly advantageous for trajectories with lower sampling efficiency, such as 3D radial imaging, as can be seen from the simulation in Figure 3b. The poorer k-space coverage compared to the cone trajectory^25^ means the radial approach benefits more from the effective sharing of information across timepoints that is achieved with a subspace reconstruction.

Compared to sliding window approaches like KWIC^20–22^, which reconstruct each frame independently with gridding, our method took advantage of the sampling across the whole train of readouts without timepoint mismatch between acquisition and reconstruction(i.e., signal from multiple TRs was combined for image of a single timepoint). Additionally, the LLR regularization applied on the subspace coefficient maps improved SNR in the final images, whereas the gridding approach used in KWIC would not provide additional denoising. Although a sliding window approach could be also combined with compress sensing to achieve additional noise suppression for high temporal resolution dynamic MRA, the computational requirements would be enormous. Compared to previously reported state-of-the-art work on 4D dynamic MRA, for which compromises had to be made either in temporal resolution^20^ (1 *mm*^3^ spatial resolution with ∼100 *ms* temporal resolution) or in spatial resolution^11^ (59 *ms* spatial resolution with 2 *mm* spatial resolution in slice direction), we achieved 1.1. *mm* isotropic spatial resolution, 14.7 *ms* temporal resolution and 200 × 200 × 125 *mm* acquisition FOV within a ∼6 *min* acquisition time. The full time-series of subspace reconstructed MRA was exhibited in Supporting Figure S3 as a dynamic image.

The proposed subspace reconstruction method could be extended to further improve the image quality of dynamic MRA. For instance, it could be combined with time-encoded PCASL to boost SNR by allowing for larger flip angle readouts and better background suppression^18^. Incorporating the kinetic model informed reconstruction into such a protocol could combine the advantages of SNR and spatiotemporal resolution for better fidelity angiograms.

While the linear compression of the signal dictionary served as a close approximation for the true underlying signal model, there are several drawbacks to this approach. Modeling the angiography signal variation of the sequence in this work is complex as the labeled blood flows in the vessels while experiencing dispersion. Additionally, the in vivo transit time varied across a wide range from close to 0 to longer than 1.8 s according to our results, which is reflected as a temporal shift in the signal timeseries. Therefore, according to our simulations, 12 principal components were required to characterize 99.95% of signal variability in the dictionary. Further increasing the number of principal components gave marginal improvement in signal fitting but increased the computational time and memory significantly. In addition, using more principal components would have weakened the prior constraint, potentially leading to increased noise sensitivity. In some extreme cases, where transit time was short or dispersion sharpness was large, the signal variation could not be represented accurately by the compressed subspace, contributing to errors in the reconstructed images. Non-linear methods like autoencoders^38^ are capable of representing complex signal variations with a smaller number of independent components (i.e., in the latent space), as demonstrated by Arefeen, et al.^39^. However, the computational convenience brought by the permutation of linear operators would be lost, and the optimization would need to be performed with a subset of frames per iteration to accommodate limited memory. The balance between computational efficiency and representation accuracy remains to be explored in greater detail in future work.

Apart from the linearly compressed representation of the kinetic model being imperfect, the model itself might not be applicable to some complex flow scenarios present in patients. For example, modeling of the complex blood flow in an aneurysm is still the subject of ongoing research^40^. Therefore, the actual generalizability of subspace representation for the kinetic model to complex scenarios requires further investigation through testing in patient cohorts.

In this preliminary study, a group of 8 subjects was scanned to prove the feasibility and advantage of the proposed model informed reconstruction for accurate qualitative and quantitative assessment. Patients with cerebrovascular disease should be recruited and studied in the future for a more comprehensive examination of the effectiveness of our method. An effort could also be made to acquire ground truth data for the validation of the accuracy of the angiography quantification results obtained from the proposed reconstruction (e.g. using a very long scan time reference).

## Conclusion

In this work, we incorporated an angiographic kinetic model into the image reconstruction process and combined this with a subspace technique to achieve ultra-high temporal resolution 4D MRA. Our method took advantage of the prior knowledge of the angiographic signal time-course to compensate for reduced sampling per frame and disentangled time-varying blood signal at different timepoints. The spatial and temporal fidelity of the resulting angiograms were, therefore, both notably improved. Parameter maps derived from the subspace results also benefited from the improved temporal fidelity and achieved better accuracy than that of the conventional temporally binned images in numerical simulation. Investigation on a group of healthy subjects demonstrated superior image quality and angiography parameter quantification using the proposed subspace reconstruction compared to the temporal binning method, consistent with the simulation results.

## CODE AND DATA AVAILABILITY STATEMENT

The code for cone trajectory used in this work could be found at

https://github.com/Michaelsqj/TrajectoryDesign.

The code for reconstruction and simulation could be found at

https://github.com/Michaelsqj/dynamic_angio.

The fabber code for model fitting could be found at

https://github.com/tomokell/fabber_models_asl/tree/capria.

We are currently unable to share subject level data due to data protection issues, although our center is actively working on a solution to this.

## Acknowledgments

We are grateful for funding support from a Sir Henry Dale Fellowship jointly funded by the Wellcome Trust and the Royal Society (220204/Z/20/Z). The Wellcome Centre for Integrative Neuroimaging is supported by core funding from the Wellcome Trust (203139/Z/16/Z) with additional support from the NIHR Oxford Health Biomedical Research Centre (NIHR203316). W.W. is supported by the Royal Academy of Engineering (RF\201819\18\92). MC is supported by the Canada Research Chairs program. The views expressed are those of the authors and not necessarily those of the NIHR or the Department of Health and Social Care. Many thanks also to Jeff Fessler, Philipp Ehses and colleagues for making available their excellent NUFFT and Siemens raw data reading MATLAB code, as well as to Siemens Healthineers for providing the base pulse sequence code, that we built upon in this work. For the purpose of open access, the author has applied a CC BY public copyright license to any Author Accepted Manuscript version arising from this submission.

## Supporting figure captions

Figure S1. Angiograms reconstructed by Extreme MRI with different regularization weights. Low regularization weight kept the fidelity of angiography but lost SNR, whereas high regularization weight overly smoothed the image and lost visibility of thin vessels, as indicated by the yellow circles. Arteries pointed by the red arrows also appeared at late timepoints only at high regularization weight, confirming these were artifacts introduced due to over-regularized on the temporal dimension.

Figure S2. Angiography reconstructed by “LLR-matched” with different regularization weights. In a similar way to Extreme MRI, the choice of regularization weight for LLR matched was a compromise between excessive smoothing and noisy images.

Figure S3. An example movie of dynamic MRA reconstructed by the subspace method shows smooth blood flow into the brain. The contrast of each frame was separately adjusted for clarity. From left to right: Maximum intensity projection (MIP) in Sagittal, coronal and transverse view.

