## Supporting Figure for "Ultra-high temporal resolution 4D angiography using arterial spin labeling with subspace reconstruction"

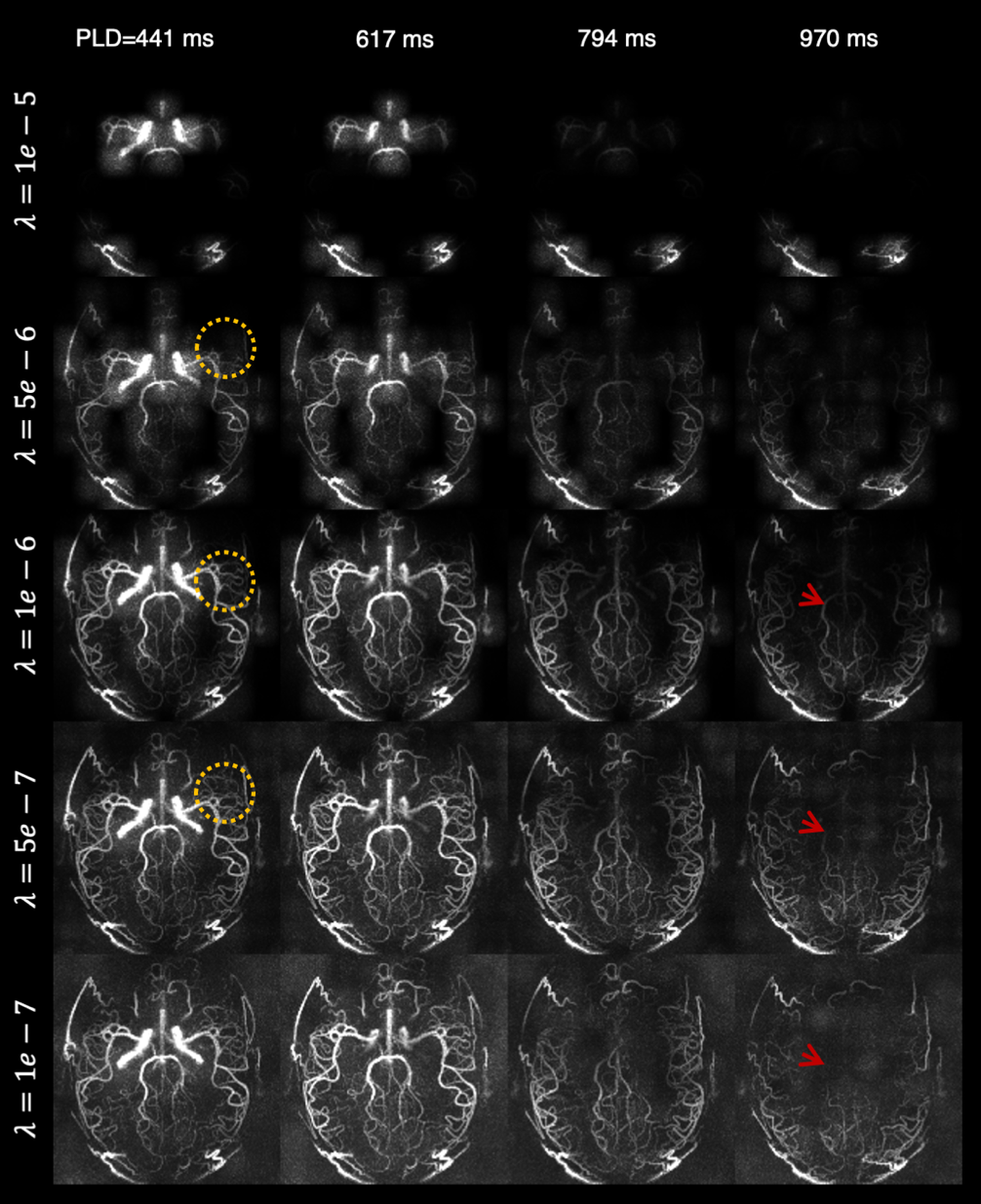

Figure S1. Angiograms reconstructed by Extreme MRI with different regularization weights. Low regularization weight kept the fidelity of angiography but lost SNR, whereas high regularization weight overly smoothed the image and lost visibility of thin vessels, as indicated by the yellow circles. Arteries pointed by the red arrows also appeared at late timepoints only at high regularization weight, confirming these were artifacts introduced due to over-regularized on the temporal dimension.

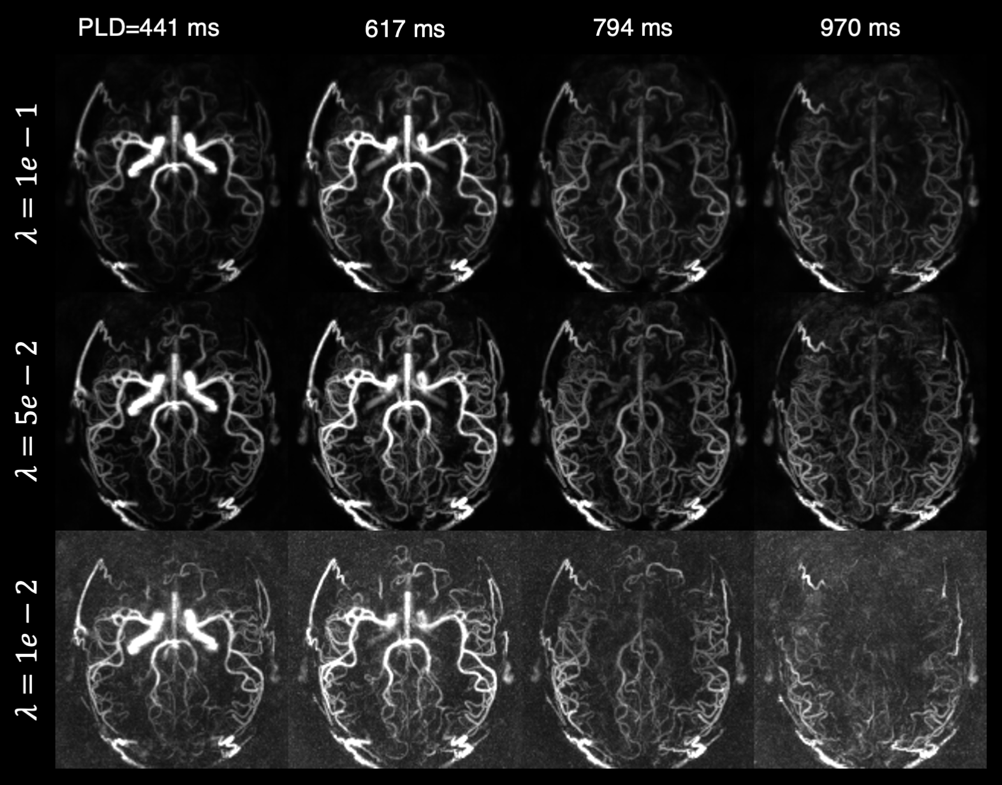

Figure S2. Angiography reconstructed by “LLR-matched” with different regularization weights. In a similar way to Extreme MRI, the choice of regularization weight for LLR matched was a compromise between excessive smoothing and noisy images.

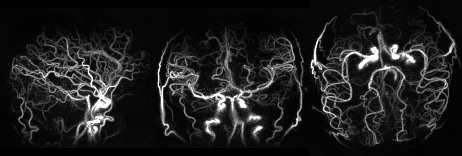

Figure S3. Example movie of dynamic MRA reconstructed by the subspace method showing smooth blood flow into the brain. The contrast of each frame was separately adjusted for clarity. From left to right: Maximum intensity projection (MIP) in Sagittal, coronal and transverse view.
